# Light-induced *psbA* translation in plants is triggered by photosystem II damage via an assembly-linked autoregulatory circuit

**DOI:** 10.1101/2020.04.27.061879

**Authors:** Prakitchai Chotewutmontri, Alice Barkan

## Abstract

The D1 reaction center protein of Photosystem II (PSII) is subject to light-induced damage. Degradation of damaged D1 and its replacement by nascent D1 are at the heart of a PSII repair cycle, without which photosynthesis is inhibited. In mature plant chloroplasts, light stimulates the recruitment of ribosomes specifically to *psbA* mRNA to provide nascent D1 for PSII repair, and also triggers a global increase in translation elongation rate. The light-induced signals that initiate these responses are unclear. We present action spectrum and genetic data indicating that the light-induced recruitment of ribosomes to *psbA* mRNA is triggered by D1 photodamage, whereas the global stimulation of translation elongation is triggered by photosynthetic electron transport. Furthermore, mutants lacking HCF136, which mediates an early step in D1 assembly, exhibit constitutively high *psbA* ribosome occupancy in the dark, and differ in this way from mutants lacking PSII for other reasons. These results, together with the recent elucidation of a thylakoid membrane complex that functions in PSII assembly, PSII repair and *psbA* translation, suggest an autoregulatory mechanism in which the light-induced degradation of D1 relieves repressive interactions between D1 and translational activators in the complex. We suggest that the presence of D1 in this complex coordinates D1 synthesis with the need for nascent D1 during both PSII biogenesis and PSII repair in plant chloroplasts.

**Significance Statement:** Photosystem II (PSII) harbors the water-splitting activity underlying oxygenic photosynthesis. The PSII reaction center protein D1 is subject to photodamage and must be replaced with nascent D1 to maintain photosynthetic activity. How new D1 synthesis is coordinated with D1 damage has been a long-standing question. Our results clarify the nature of the light-induced signal that activates D1 synthesis for PSII repair in plants, and suggest an autoregulatory mechanism in which degradation of damaged D1 relieves a repressive interaction between D1 and translational activators in a complex that functions in PSII assembly and repair. This proposed mechanism comprises a responsive switch that couples D1 synthesis to need for D1 during PSII biogenesis and repair.

## Introduction

Oxygenic photosynthesis shapes earth’s ecosystems by generating molecular oxygen, consuming atmospheric CO_2,_ and producing the carbohydrates that fuel life. Oxygenic photosynthesis is mediated by a set of protein-prosthetic group complexes embedded in the thylakoid membranes of cyanobacteria and chloroplasts: Photosystem I (PSI), Photosystem II (PSII), the cytochrome *b*_*6*_*f* complex (cyt *b*_*6*_*f*), and ATP synthase (1,2). PSII drives this process by using the energy of sunlight to extract electrons from water, producing molecular oxygen as a byproduct (3,4). However, exposure to light also damages the PSII reaction center. An elaborate repair cycle involves the partial disassembly of PSII, selective proteolysis of D1, its replacement with newly synthesized D1, and reassembly of PSII (5-7). D1 is damaged even at low light intensities, but the rate of damage increases with light intensity (8-10). D1 synthesis keeps pace with this damage up to moderate light intensities; beyond this point, D1 synthesis plateaus or even decreases, and photosynthesis is inhibited (8,9,11). In chloroplasts, the PSII repair cycle is promoted by membrane rearrangements that expose the damaged core to proteases that degrade D1 and to the ribosomes that synthesize new D1 (6,7,12).

The PSII repair cycle has come under scrutiny due to its importance for maintaining photosynthesis. Considerable progress has been made in understanding the source of D1 damage, the architectural reorganization of thylakoid membranes, the disassembly of damaged PSII, the proteases that degrade damaged D1, and factors that promote PSII reassembly (5-7,12). However, mechanisms underlying the selective translation of *psbA* mRNA to provide D1 for PSII repair remain obscure (13,14). The consensus view in recent years has been that *psbA* translation for PSII repair is regulated at the elongation step (7,15-17), a view that arises primarily from experiments with the green alga *Chlamydomonas reinhardtii* (Chlamydomonas) (18). However, we showed recently that regulated translation initiation makes a large contribution in plants (19). These experiments used ribosome profiling (ribo-seq) to monitor ribosome occupancy on chloroplast open reading frames (ORFs) in maize and Arabidopsis upon shifting seedlings harboring mature chloroplasts between light and dark. The results showed that ribosome occupancy on *psbA* mRNA increases dramatically within fifteen minutes of shifting plants from the dark to the light and drops rapidly after shifting plants from light to dark. Furthermore, *psbA* mRNA is the only chloroplast mRNA to exhibit a substantive change in ribosome occupancy following short term light-dark shifts, and this results in an “over-production” of D1 (with respect to other PSII subunits) in the light. These observations imply that the light-induced recruitment of ribosomes to *psbA* RNA in mature plant chloroplasts serves the purpose of PSII repair. The same study revealed a plastome-wide increase or decrease in translation elongation rate following a shift to light or dark, respectively. Thus, although the rate of translation elongation on *psbA* mRNA decreased in the dark, it remains unclear whether a *psbA*-specific change in translation elongation rate contributes to the control of D1 synthesis for PSII repair in plants.

A related issue concerns the nature of the light-induced signal that triggers D1 synthesis for PSII repair. The prevailing view in recent years is that light induces D1 synthesis via its effects on products of photosynthetic electron transport (7,15-17). Proposed signals include ATP, the *trans*-thylakoid proton gradient, reduced plastoquinone, and dithiol reduction (20-26). However, the experiments that led to this view did not distinguish plastome-wide changes in translation rate from the *psbA*-specific activation that provides D1 for PSII repair. These two phenomena may involve distinct signals, so the parsing of global effects from *psbA*-specific effects is essential for elucidating the signals.

In this study, we sought to clarify the light-induced signal that triggers the recruitment of ribosomes to *psbA* mRNA in mature plant chloroplasts. For this purpose, we analyzed light-induced D1 synthesis and *psbA* ribosome occupancy in response to different wave lengths of light, and in mutants lacking various components of the photosynthetic apparatus. Our results show that the light-activated recruitment of ribosomes to *psbA* mRNA is not mediated by the activation of photosynthetic electron transport. Instead, our results strongly suggest that light triggers this response by causing D1 damage, a hypothesis that had been proposed in early studies (27,28) but that has lost prominence. We show further that the absence of the PSII assembly factor HCF136, which promotes a very early step in D1 assembly (29-32), results in constitutively high *psbA* ribosome occupancy in the dark. These results reveal an intimate connection between a specific early PSII assembly intermediate, light-induced D1 damage, and light-regulated *psbA* ribosome occupancy. We propose that the recently elucidated HCF244/OHP1/OHP2 complex, which functions in both PSII assembly and *psbA* translational activation (32-35), serves as the hub of an autoregulatory mechanism that links D1 synthesis to need for nascent D1 during PSII repair and biogenesis.

## Results

### Action Spectrum Analysis Supports D1 Damage as the Light-Induced Trigger for *psbA* Translation in Mature Chloroplasts

It is widely believed that light induces *psbA* translation via its effects on intermediates or products of photosynthesis (reviewed by 7,15,16,17,36). However, early reports proposed instead that D1 synthesis is induced by light-induced D1 degradation (27,28). Furthermore, the existence of both plastome-wide and *psbA-*specific effects of light on chloroplast translation (19) complicates interpretation of much of the prior research on this topic, as these two effects may involve distinct signals.

To distinguish between the two proposed signaling modes, we exploited the fact that the action spectrum for PSII damage differs from that for photosynthesis (11,37-40). In particular, UV-A is particularly potent at inducing D1 damage but is an inefficient driver of photosynthesis. This effect of UV-A light has been attributed to its absorption by the manganese cluster associated with the PSII reaction center (reviewed in 41). If light-induced D1 damage is the triggering event for *psbA* ribosome recruitment, then the action spectrum for *psbA* ribosome recruitment would match that for D1 damage and not that of photosynthesis.

To address this possibility, we used the scheme shown in Figure 1*A*. Arabidopsis seedlings were grown in moderate light under diurnal cycles for several weeks. They were then acclimated to very low intensity light (8 µmol m^−2^ s^−1^) for 30 minutes to minimize photodamage. Supplemental low-intensity illumination was then provided with either white light (25 µmol m^−2^s^−1^) or with monochromatic UV-A, blue, green, or red light (5 µmol m^−2^s^−1^). The red light (635 nm) is in the peak of the photosynthetically-active region of the spectrum, the green and blue lights (522 nm and 469 nm, respectively) are roughly two-fold less efficient at driving photosynthesis, and the UV-A light (370 nm) is an additional several fold less efficient (40). Plants were harvested fifteen minutes after exposure to the supplemental light for analysis by ribosome profiling, or were subjected to *in vivo* pulse labeling during an additional 20 minutes under the same supplemented light condition. The pulse-labeling assays were performed in the presence of cycloheximide to inhibit cytosolic protein synthesis.

**Fig. 1.**
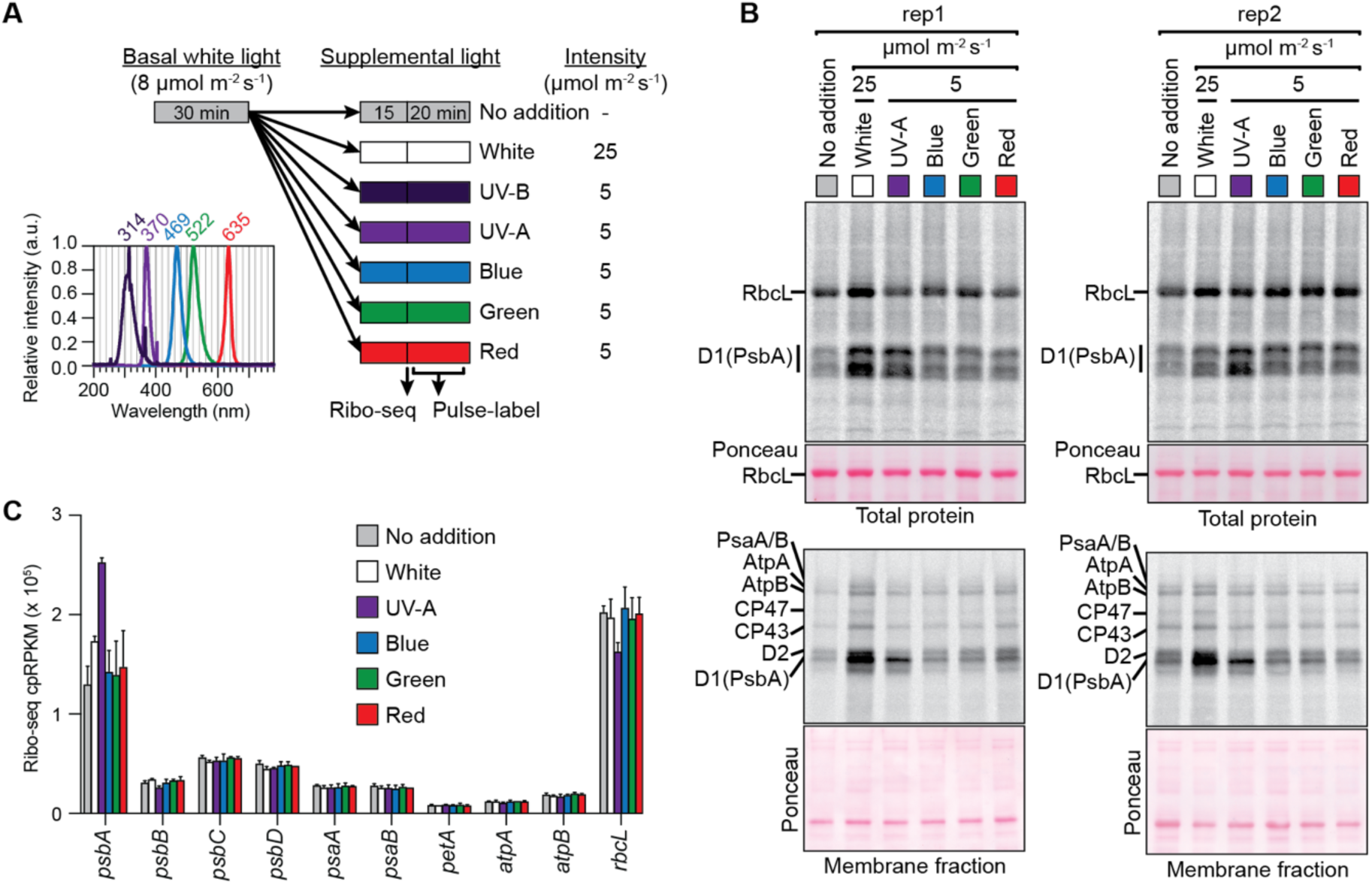
Action spectrum analysis of light-induced *psbA* translation in Arabidopsis. (*A*) Light treatment regime and spectra of each monochromatic light source. Plants were acclimated to low intensity (8 µmol m^−2^ s^−1^) white light for 30 min prior to transfer to the indicated supplemental light. Fifteen minutes later, plants were harvested for ribo-seq or were used for pulse labeling. Samples were loaded on the basis of equal chlorophyll. (*B*) Pulse labeling analysis of plants exposed to light regimes diagrammed in (*A)*. Excised aerial portions of seedlings were radiolabeled for 20 min in the presence of cycloheximide. Total leaf lysates (top) and membrane fractions (bottom) were resolved by SDS-PAGE, transferred to nitrocellulose, and radiolabeled proteins were detected by phosphorimaging. The chloroplast-encoded proteins RbcL, D1 (PsbA), D2 (PsbD), CP47 (PsbB), CP43 (PsbC), AtpA, AtpB, and PsaA/B are marked. The nitrocellulose filters with the bound radiolabeled proteins were stained with Ponceau S (below) to illustrate relative sample loading. (*C*) Normalized ribosome footprint abundance for chloroplast genes following the indicated light treatment. The mean ± SD (from two replicates) is shown. The values for all chloroplast genes are available in *SI Appendix, Dataset S1*.

The pulse-labeling data showed that supplemental UV-A light caused a greater increase in the rate of D1 synthesis than did any of the other single wavelengths sampled (Fig. 1*B*). For example, analysis of total leaf lysates (upper panels) showed that the labeling of D1 in comparison to the soluble protein RbcL is elevated by the UV-A treatment, but not by the other treatments. Similarly, analysis of the membrane fraction (bottom panels) showed that the labeling of D1 in comparison to other membrane proteins was most strongly increased by UV-A supplementation. Analysis of the same material by ribosome profiling (Fig. 1*C* and *SI Appendix Dataset S1*) showed that the supplemental UV-A light caused a specific increase in ribosome occupancy on *psbA* mRNA (roughly two-fold) whereas the green, blue and red lights had no significant effect. The fact that UV-A more effectively stimulated a specific increase in D1 synthesis and *psbA* ribosome recruitment than did photosynthetically-active wavelengths argues against models that invoke products of photosynthesis as triggers for the *psbA-*specific response, and is consistent with the possibility that the trigger is light-induced D1 damage.

Activation by UV-A could potentially be mediated by one of the photoreceptors known to absorb UV-A: cryptochromes, phototropins, or UVR8 (42). However, this possibility seems unlikely for several reasons. First, peak absorption for phototropins and crytochromes is in the blue region of the spectrum (43,44), which did not trigger *psbA* translation. Second, these photoreceptors do not localize to chloroplasts, making them unlikely transducers of the very rapid *psbA* translation response. Nonetheless, we investigated the potential involvement of UVR8, whose peak response is in the UV-B (280-315 nm) (45). Supplementation of low intensity white light with a low dose of UV-B inhibited synthesis of all detected proteins in wild-type plants (Fig. 2*A*), implying a global inhibition of chloroplast translation. Furthermore, pulse-labeling analysis of a *uvr8* mutant showed normal light-induced D1 synthesis (Fig. 2*B*).

**Fig. 2.**
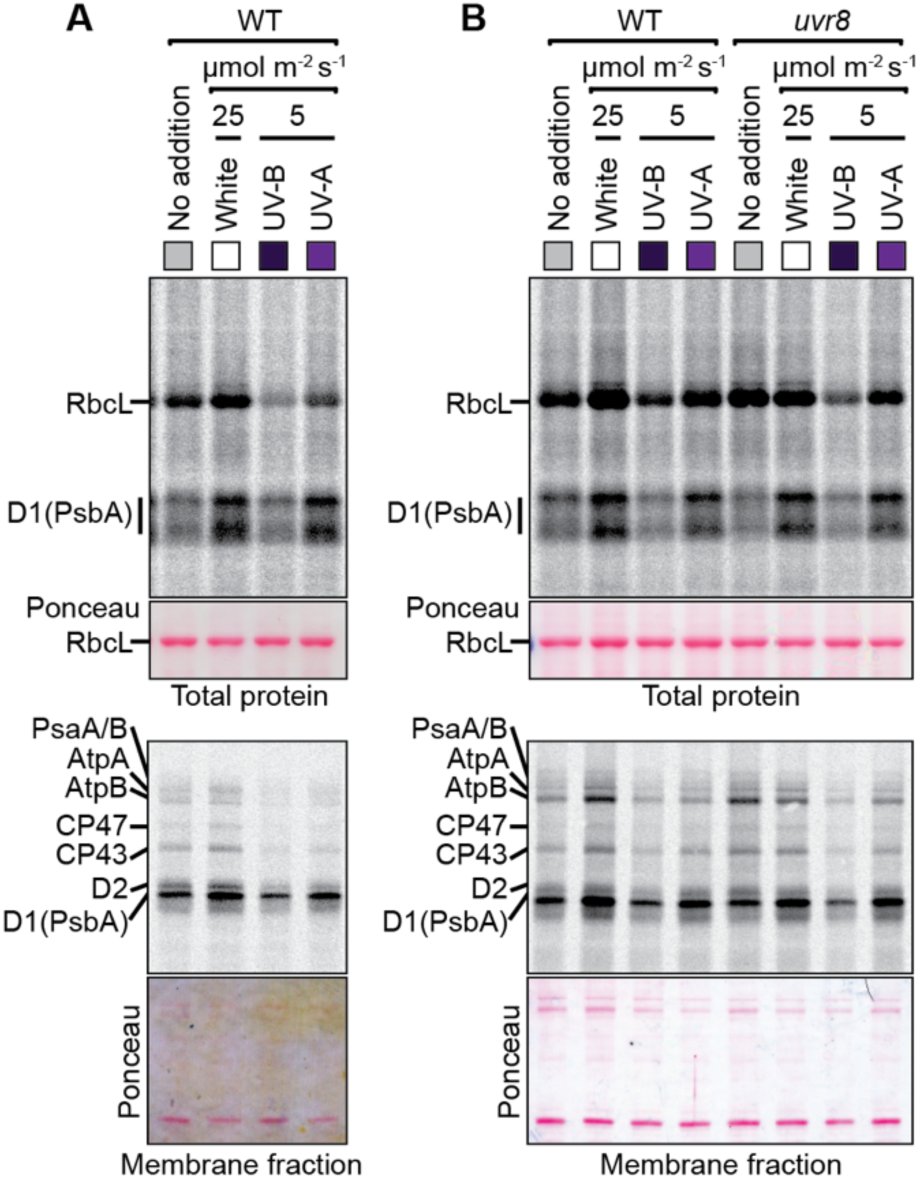
Pulse labeling analysis of the effects of UV-B on chloroplast translation. The experiment was performed as in Fig. 1*A*, with supplemental light provided by either UV-B or UV-A as indicated. Excised aerial portions of seedlings were radiolabeled for 20 min in the presence of cycloheximide. Total leaf lysates (top) and membrane fractions (bottom) were resolved by SDS-PAGE and transferred to nitrocellulose, and radiolabeled proteins were detected by phosphorimaging. Samples were loaded on the basis of equal chlorophyll. The nitrocellulose filters with the bound radiolabeled proteins were stained with Ponceau S (below) to illustrate relative sample loading. *(A)* Analysis of 14-day old wild-type (WT) plants. *(B)* Analysis of young rosette leaves from 55-day-old wild-type or *uvr8* mutant plants.

The findings above strongly suggest that the *psbA-*specific light response – whether assayed by pulse-labeling or ribosome profiling – is triggered by light-induced D1 damage and not by products or intermediates of photosynthetic electron transport. In addition, two aspects of the data revealed that a different signal triggers the plastome-wide effects on translation elongation rate. First, the supplemental white light increased synthesis of all proteins detected by pulse labeling (Fig. 1*B*) with no corresponding change in ribosome occupancy on their mRNAs (except *psbA*) (Fig. 1*C*). Second, the supplemental white light resulted in greater D1 synthesis but less increase in *psbA* ribosome occupancy than did the weaker UV-A light (Fig. 1*B*,*C*). These results suggest that the white light increased synthesis of many (if not all) chloroplast-encoded proteins primarily through a concerted increase in elongation and initiation rate that did not change ribosome occupancy, whereas the UV-A treatment acted specifically on *psbA* at the level of ribosome recruitment. Thus, light-induced recruitment of ribosomes to *psbA* mRNA is likely triggered by D1 damage, whereas the plastome-wide increase in translation elongation rate may be triggered by a product of photosynthesis.

### Normal Light-Induced D1 Synthesis in Mutants Lacking the Chloroplast ATP Synthase, PSI, or the STN7/STN8 Protein Kinases

In a complementary approach, we addressed the role of photosynthesis in light-activated chloroplast translation by using maize and Arabidopsis mutants lacking specific components of the photosynthetic apparatus. To minimize energy deficits resulting from the photosynthetic defects, Arabidopsis mutants were grown on sucrose-containing medium and maize mutants were analyzed prior to the depletion of seed reserves. Non-photosynthetic maize mutants are similar in size and morphology to their normal siblings at this stage (see, e.g. *SI Appendix, Fig. S1A* and (33)), whereas non-photosynthetic Arabidopsis mutants show varying degrees of developmental delay despite the sucrose supplementation (see, e.g. (46-48). Plants were grown in diurnal cycles, and translation was assayed by pulse-labeling at midday, after one hour in the dark, or following 15 minutes of reillumination (Fig. 3).

**Fig. 3.**
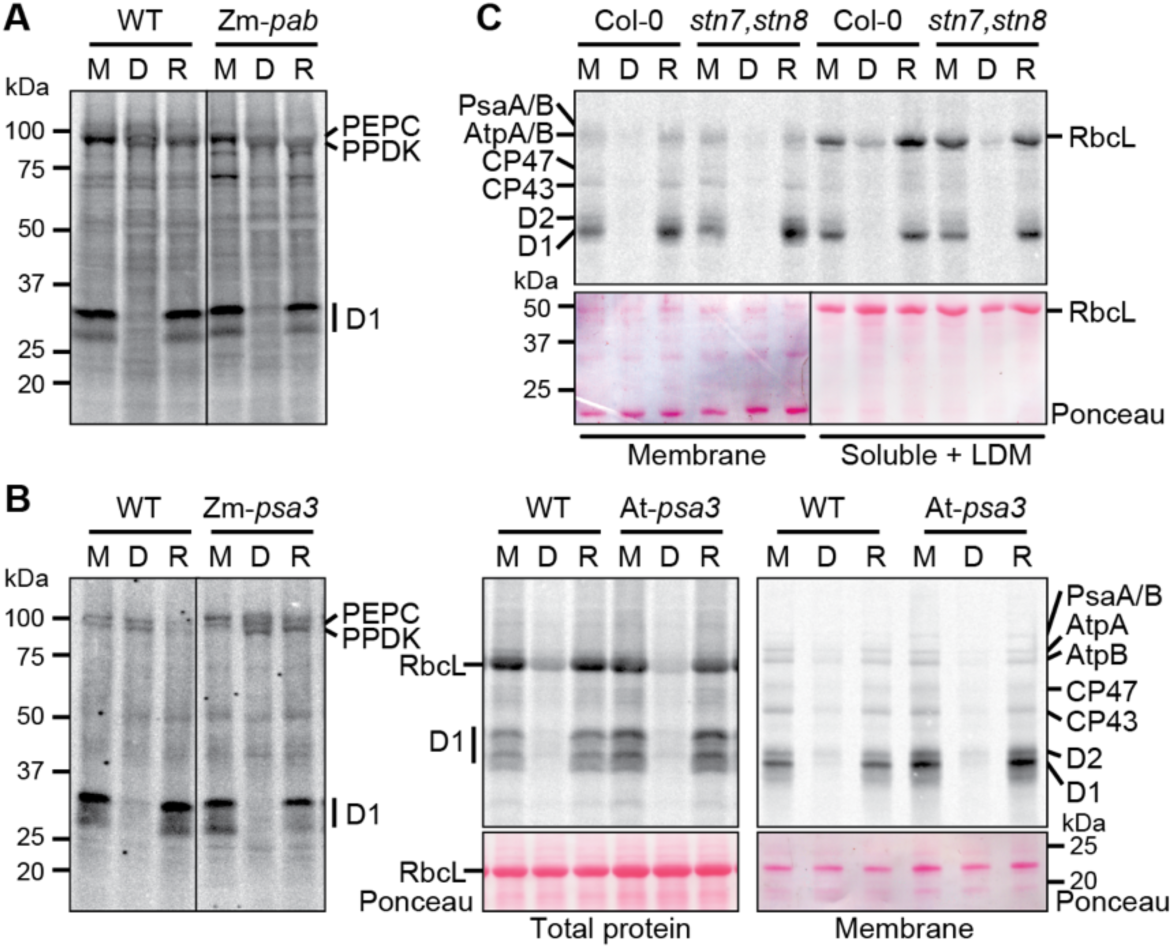
Analysis of light-regulated D1 synthesis in mutants lacking ATP synthase, PSI, or the STN7 and STN8 kinases. Plants were pulse-labeled at midday (M), after 1-h in the dark (D), or after 15-min reillumination (R). Maize mutants (Zm-*pab* and Zm-*psa3)* were radiolabeled for 15 min in the absence of cycloheximide, and total leaf lysates were analyzed by SDS-PAGE with sample loading normalized based on labeling of two cytosolic proteins at ∼100 kDa (presumed to be PEPC and PPDK). Arabidopsis mutants (At-*psa3* and *stn7,stn8*) were radiolabeled for 20 min in the presence of cycloheximide, and samples were loaded on the basis of equal chlorophyll. (*A*) Analysis of a maize Zm-*pab* mutant, lacking the thylakoid ATP synthase. The mutant is described in *SI Appendix, Fig. S1A*. Total leaf lysates are shown. The line separates non-contiguous lanes from the same exposure of the same gel. (*B*) Analysis of mutants lacking PSI in maize (Zm-*psa3*) and Arabidopsis (At-*psa3*). Arabidopsis samples were analyzed both as total leaf lysates (left) and as membrane fractions (right). The nitrocellulose filters harboring the radiolabeled proteins were stained with Ponceau S to illustrate equal sample loading. RbcL-large subunit of Rubisco. The line in the Zm-*psa3* panel separates non-contiguous lanes from the same exposure of the same gel. (*C*) Analysis of an Arabidopsis *stn7,stn8* double mutant. The experiment was performed as for At-*psa3* except that the soluble fraction also included low-density membranes (LDM). A single Ponceau S-stained blot was imaged with different contrast (marked by a line) to improve visibility of the bands.

We examined the role of the chloroplast ATP synthase by analysis of a maize mutant lacking the ortholog of Arabidopsis PAB, which is required for ATP synthase assembly (49). Immunoblot analyses of the maize mutant (Zm-*pab*) showed a roughly 20-fold decrease in the abundance of the AtpB subunit of the ATP synthase, with no impact on the abundance of core subunits of other photosynthetic complexes (*SI Appendix, Fig. S1A*). The Zm-*pab* mutant showed robust light-induced D1 synthesis (Fig. 3*A*). These results suggest that light-induced ATP synthesis is not required to trigger D1 synthesis when sufficient ATP is available from other sources to support the energy demands of translation.

We examined the need for PSI by analysis of *psa3* mutants, which lack PSI due to a defect in PSI assembly (50) (*SI Appendix, Fig. S1B*). The *psa3* mutants in both maize and Arabidopsis showed robust light-induced D1 synthesis (Fig. 3*B*), suggesting that processes that require PSI are not required to trigger D1 synthesis. In fact, the Arabidopsis mutant showed elevated D1 synthesis under moderate light intensity, an effect that was reduced under very low intensity light (*SI Appendix, Fig S1C*). This correlates with the increased susceptibility of PSII to light-induced damage in Arabidopsis mutants lacking PSI (51,52), consistent with the hypothesis that D1 damage triggers the *psbA-*specific response.

Two paralogous protein kinases, STN7 and STN8, are involved in the adaptation of the photosynthetic apparatus to unbalanced photosystem excitation or excess light energy (reviewed in 53). STN7 phosphorylates light-harvesting chlorophyll *a/b* binding proteins and is required for state transitions (54), whereas STN8 phosphorylates PSII core proteins and promotes PSII repair (53). To address whether STN7 and STN8 contribute to the regulation of D1 synthesis in response to light, we pulse-labeled an Arabidopsis *stn7-stn8* double mutant (55) in light and dark (Fig. 3*C*). Light-induced D1 synthesis was not altered in the double mutant, indicating that the STN7 and STN8 kinases are not necessary for the signal transduction cascade that triggers light-induced D1 synthesis in mature chloroplasts.

### Pharmacological and Genetic Data Support the View that Light Induces the Plastome-Wide Translation Response via Electron Transport-Induced Effects on pH

Experiments involving pharmacological treatments of lysed barley chloroplasts suggested that light induces a general increase in translation elongation rate via electron-transport induced changes in pH (21). In accord with that view, the PSII inhibitor DCMU caused a general inhibition of translation in isolated Chlamydomonas chloroplasts (20), in Chlamydomonas cells (56), and in cyanobacteria (57). Those observations suggest that photosynthetic electron transport underlies the plastome-wide effects of light we detected by ribo-seq (19), a possibility that is also supported by the action spectrum data presented above.

To examine this issue further, we used pulse-labeling to analyze chloroplast protein synthesis in mutants lacking the cyt *b*_*6*_*f* complex (Fig. 4*A*). The cyt *b*_*6*_*f* complex transports electrons from PSII to PSI and pumps protons from the stroma to the thylakoid lumen. We analyzed orthologous mutants in maize and Arabidopsis, *pet2* and *hcf208* respectively, which lack the cyt *b*_*6*_*f* complex due to a defect in heme attachment (47,58). Light-induced D1 synthesis in the maize *pet2* mutant was undetectable (Fig. 4*A*, left panel). However, D1 was the only chloroplast-encoded protein whose synthesis we could reliably detect in maize, so we could not discern whether this was a *psbA-*specific or plastome-wide defect. The Arabidopsis *hcf208* mutant also showed a considerable reduction in D1 synthesis in the light (Fig. 4*A*, right panel). This was particularly apparent when the membrane fractions were analyzed on gels containing urea (right panel), which resolve the closely-migrating D1 and D2 bands. Furthermore, synthesis of all of the detected membrane proteins was reduced in the *hcf208* mutant, as was synthesis of the soluble protein RbcL. This behavior suggests that the cyt *b*_*6*_*f* complex is required for the plastome-wide activation of translation elongation by light.

**Fig. 4.**
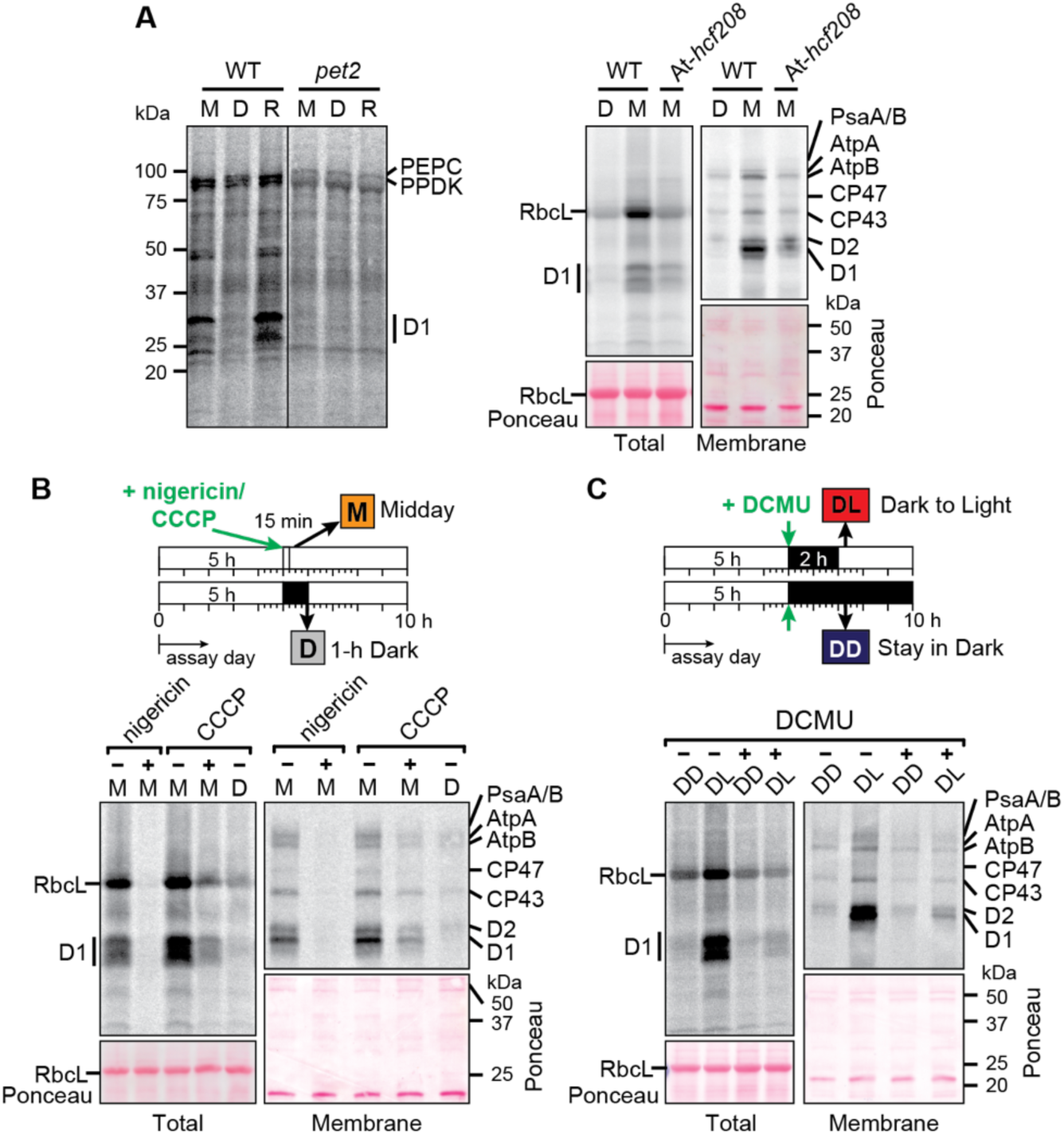
Analysis of light-regulated chloroplast protein synthesis in mutants lacking the cytochrome *b*_*6*_*f* complex or after treatment with nigericin, CCCP or DCMU. (*A*) Analysis of cytochrome *b*_*6*_*f* mutants. Pulse-labeling was performed at midday (M), after 1-h in the dark (D), or after 15-min reillumination (R). The left panel shows total leaf lysates of the maize *pet2* mutant labeled in the absence of cycloheximide. The line separates non-contiguous lanes on the same gel. The two rapidly synthesized proteins at ∼100 kDa are likely to be the nuclear gene products PEP Carboxylase and PPDK. The right panel shows total leaf lysates and membrane fractions of Arabidopsis At-*hcf208* mutants labeled in the presence of cycloheximide. The Ponceau S-stained blots show relative sample loading. Samples were loaded on the basis of equal total protein. (*B*) Arabidopsis seedlings (Col-0) were treated with 25 µM nigericin or 100 µM CCCP for 15 min before pulse-labeling (20 min) in the presence of cycloheximide in the light at midday (M). Mock treated plants were radiolabeled after 1-h in the dark (D) for comparison. Samples were loaded on the basis of equal chlorophyll. (*C*) Arabidopsis seedlings (Col-0) were treated with 20 µM DCMU in darkness for 2 h before shifting to light for 15 min (DL) or remaining in the dark (DD), followed by 20-min labeling in the presence of cycloheximide. Samples were loaded on the basis of equal chlorophyll.

Loss of the cyt *b*_*6*_*f* complex would affect numerous potential signals for the plastome-wide translational response, including ATP, reduced thioredoxin, plastoquinone redox state, and the *trans*-thylakoid proton gradient. However, the fact that light-induced chloroplast protein synthesis was normal in PSI and ATP synthase mutants (Fig. 3) suggests that ATP and reduced thioredoxin are not relevant. To distinguish between the remaining possibilities, we analyzed effects of several pharmacological inhibitors of photosynthesis. Treatment of Arabidopsis with the uncouplers nigericin or CCCP inhibited the synthesis of all detected plastid-encoded proteins in the light (Fig. 4*B*), consistent with a role for light-induced changes in pH or the *trans*-thylakoid proton motive force in the signaling pathway. Treatment with DCMU in the dark phase also inhibited light-induced chloroplast translation (Fig. 4*C*), similar to effects reported previously in Chlamydomonas (20,56). The fact that DCMU and the absence of the cyt *b*_*6*_*f* complex had similar inhibitory effects on translation in the light but have opposite effects on the redox state of the plastoquinone pool suggest that plastoquinone redox state is not relevant to the plastome-wide translational induction by light. Taken together, our inhibitor and genetic data support the prior conclusion from the lysed chloroplast system (21) that light-induced effects on pH trigger a global increase in chloroplast translation elongation rate.

### Constitutive *psbA* Ribosome Occupancy in *hcf136* Mutants Implicates an Early PSII Assembly Intermediate as a Repressor of *psbA* Translation in the Dark

We next used ribo-seq to examine the role of PSII in the light-induced recruitment of ribosomes specifically to *psbA* mRNA. In one experiment, Arabidopsis seedlings were treated with DCMU using the same regime employed for the pulse-labeling assay (Fig. 3*C*), and were then harvested for ribosome profiling during the dark phase or after 15 minutes of reillumination (Fig. 5*A*). DCMU reduced the recruitment of ribosomes to *psbA* mRNA in response to light and did not affect ribosome occupancy on other ORFs (Fig. 5*A* and *SI Appendix, Fig. S2*). Notably, however, the degree to which DCMU inhibited light-induced *psbA* ribosome recruitment was modest in comparison with its strong effect on D1 synthesis in the pulse labeling assay (Fig. 4*C*). These results suggest that DCMU inhibits light-induced D1 synthesis in two ways: (i) by interfering with the light-induced recruitment of ribosomes specifically to *psbA* mRNA, and (ii) by reducing translation elongation rate on *psbA* and other chloroplast mRNAs.

**Fig. 5.**
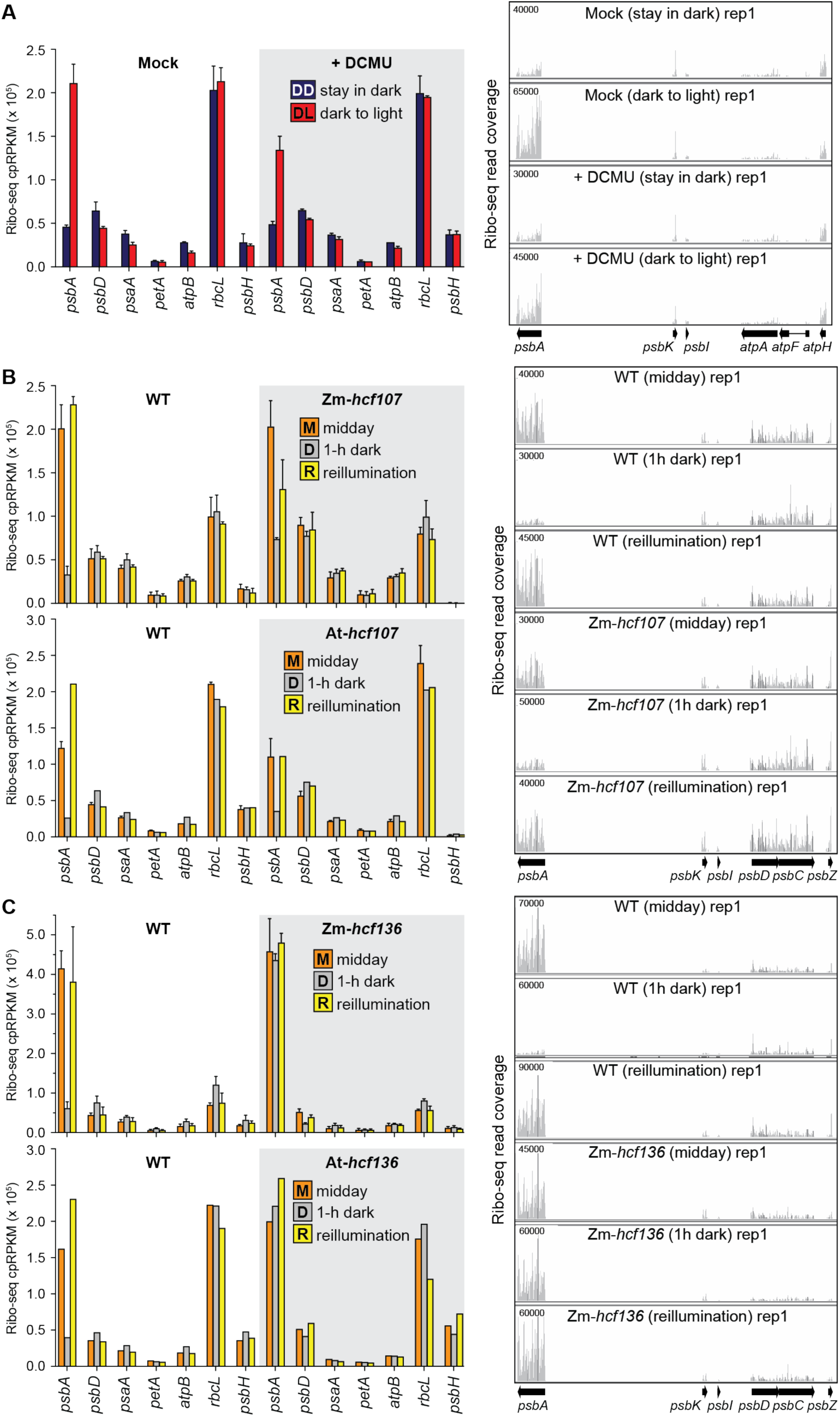
Ribo-seq analysis of effects of PSII disruptions on chloroplast ribosome occupancies in dark and light. Left panels show ribosome footprint abundance for *psbA* and several other chloroplast genes. The values for all chloroplast genes are available in *SI Appendix, Dataset S1*. Mean ± SD is shown for experiments that were performed with replicates. Several of the Arabidopsis mutant analyses lacked replicates, but results from the orthologous maize mutants demonstrate replicability. Right panels show screen captures from the Integrated Genome Viewer (IGV) that display ribosome footprint coverage along *psbA* and adjacent ORFs. Y-axis values indicate the number of reads at each position (not normalized). The maximum Y-axis values (shown in upper left) were chosen such that the magnitude for ORFs adjacent to *psbA* were similar among the conditions being compared. (*A*) Analysis of Arabidopsis (Col-0) after DCMU treatment. The treatments were as described in Fig. 4*C* and were performed in two replicates. IGV screen captures of the second replicate are shown in *SI Appendix, Fig. S2*. (*B*) Analysis of Zm-*hcf107* and At-*hcf107* mutants. The data show the known loss of *psbH* expression in these mutants (60,81). Two replicates were performed for Zm-*hcf107*. One of the replicates of the midday At-*hcf10*7 samples was published previously (81). The IGV images are from maize data. IGV screen captures of replicates are shown in *SI Appendix, Fig. S3*. (*C*) Ribo-seq analysis of Zm-*hcf136* and At-*hcf136* mutants. The midday data were published previously (33) and are shown here again in the context of the complete experiment. The Zm-*hcf136* experiment was performed with two replicates. The IGV images are from maize data. IGV captures of replicates, and chloroplast mRNA abundance (RNA-seq data) are shown in *SI Appendix, Fig. S4*.

In a second approach, we examined light-induced *psbA* ribosome recruitment in two mutants lacking PSII: *hcf107*, which lacks PSII due to a defect in expressing the chloroplast *psbH* gene (59-61), and *hcf136*, which lacks PSII due to a defect at an early step in PSII assembly (29-31). In both cases, we analyzed orthologous mutants in maize and Arabidopsis, with leaves harvested at midday, after one hour in the dark, and after 15 minutes of reillumination. Light-induced changes in *psbA* ribosome occupancy were fairly normal in the *hcf107* mutants (Fig. 5*B* and *SI Appendix, Fig. S3*), although the response to reillumination may be muted. However, the response in *hcf136* mutants was dramatically altered (Fig. 5*C* and *SI Appendix, Fig. S4*): *psbA* ribosome occupancy was normal (or perhaps elevated) in the light and did not decrease in the dark. This striking behavior indicates that HCF136 is required to repress the recruitment of ribosomes to *psbA* mRNA in the dark, thereby linking PSII assembly to light-regulated *psbA* translation.

This effect of HCF136 is particularly intriguing in light of the functional relationship between HCF136 and a recently elucidated complex harboring proteins denoted HCF244, OHP1, and OHP2 (32,34,35,62,63). This complex and its cyanobacterial ortholog (Ycf39/HliD/HliC) binds nascent D1 and is required to assemble D1 with other reaction center proteins (32,34,63). Current data suggest that HCF136 and its cyanobacterial ortholog (Ycf48) act upstream of the HCF244 complex during PSII assembly, and are required to incorporate nascent D1 into the HCF244 complex (29-31,64,65). HCF136 and the HCF244/OHP1/OHP2 complex are involved in both PSII assembly and PSII repair (30,32,63). Furthermore, the HCF244 complex in plants is also required for *psbA* translation initiation (33,46), providing a molecular link between D1 assembly status and *psbA* translation.

These recent insights into the HCF244 complex, in conjunction with evidence that unassembled D1 represses *psbA* translation in Chlamydomonas (18,66), underlie our recent proposal that nascent D1 negatively autoregulates *psbA* translation through inhibitory interactions with HCF244/OHP1/OHP2. The constitutively high *psbA* ribosome occupancy in *hcf136* mutants is consistent with this scheme: *hcf136* mutants cannot incorporate D1 into the HCF244 complex, leaving the translation activation function of the complex constitutively “on”. These *hcf136* data allow us to expand the scope of the HCF244/OHP1/OHP2-centered translational autoregulatory model to incorporate light-regulated *psbA* translation and PSII repair, as discussed below.

## Discussion

The experiments described here address two long-standing questions in plant biology: how the rate of D1 synthesis is coupled to need for nascent D1 following PSII photodamage, and how *psbA* translation is activated in response to light. Our results show that these two phenomena are intimately connected. Our data strongly suggest that light induces *psbA* translation in mature chloroplasts by triggering D1 damage, and not by affecting metabolites of photosynthesis as has often been assumed. Our findings also link high *psbA* ribosome occupancy with the absence of D1 from a complex that is required for PSII assembly, PSII repair and *psbA* translation. These results, together with prior literature, suggest an autoregulatory mechanism in which the presence of D1 in the assembly/repair complex negatively regulates D1 synthesis via inhibitory interactions with *psbA* translational activators in the complex. According to this model, light-induced D1 degradation relieves this repression, thereby triggering the recruitment of ribosomes to *psbA* mRNA. This model and the evidence for it are elaborated below.

### Distinct Light-Induced Signals Trigger the Plastome-Wide and *psbA-*Specific Translation Response

When leaves harboring mature chloroplasts are shifted from dark to moderate light, chloroplast translation is rapidly activated in two ways: the recruitment of ribosomes specifically to *psbA* mRNA is superimposed on a plastome-wide increase in elongation rate that does not change ribosome occupancy on other ORFs (19). Results presented here demonstrate that light triggers these two responses via different signals. Our data show that the *psbA-*specific response does not require photosynthetic electron transport. For example, UV-A was more effective than photosynthetically-active wavelengths at triggering D1 synthesis and *psbA* ribosome recruitment, and these effects were specific for D1/*psbA*. Furthermore, light-induced D1 synthesis was not disrupted in mutants with severe PSI or ATP synthase deficiencies. By contrast, the synthesis of all detected chloroplast-encoded proteins was reduced by inhibitors of PSII or photophosphorylation and in a mutant lacking the cyt *b*_*6*_*f* complex. Taken together, these data indicate that the plastome-wide response is triggered by a product of photosynthetic electron transport, whereas the *psbA-*specific response is not.

A study with lysed barley chloroplasts concluded that light-induced changes in pH globally activate translation elongation (21), and our *in vivo* data support that view. However, our findings run counter to the widespread view that light specifically activates D1 synthesis via its effects on photosynthetic electron transport (reviewed by 7,15,16,17). The evidence for this view comes primarily from the use of chemical inhibitors. Although those treatments inhibited D1 synthesis, our results and prior studies (21,56) indicate that such treatments globally repress chloroplast translation. Misdirection on this issue may have arisen from the fact that some studies only examined D1 synthesis, and others highlighted effects on D1 even though general inhibition was apparent (e.g. 20,22). Limiting ATP has also been proposed to be a key control point for D1 repair synthesis {Murata, 2018 #131,25). Although ATP is a critical resource for translation, our results show that photophosphorylation is not an activating signal for *psbA* ribosome recruitment and D1 repair synthesis in mature chloroplasts.

### Light-Induced D1 Damage as the Trigger for *psbA* Translation in Mature Chloroplasts

D1 damage occurs at all light intensities, but photoinhibition occurs only when the rate of damage exceeds the rate of PSII repair (8-11). Thus, light-induced D1 synthesis for PSII biogenesis *versus* PSII repair cannot be distinguished simply on the basis of light intensity, as is often assumed in discussions of this topic. Although we used moderate “growth” light intensities, we infer that the bulk of the light-induced D1 synthesis in our experiments represents repair synthesis because (i) we analyzed leaf tissue harboring mature chloroplasts that had completed the biogenesis phase, and (ii) D1 is over-produced with respect to other PSII subunits at this stage, whereas D1 is produced stoichiometrically with other PSII subunits in immature chloroplasts (67).

The most parsimonious explanation of our data is that light induces D1 synthesis in mature chloroplasts by triggering PSII damage. This view harkens back to mechanisms that had been suggested in early studies (27,28,68), but that had lost visibility in recent years (7,15,16,69). A key piece of evidence we present for this conclusion is that UV-A light more efficiently stimulated D1 synthesis and *psbA* ribosome recruitment than did red, green or blue light (Fig. 1), correlating with the action spectrum of PSII damage and not with that of photosynthesis (11,37-40). An alternative scenario could involve a UV-absorbing photoreceptor, but this is unlikely for several reasons. First, light induces D1 translation in isolated chloroplasts (reviewed in 14) but receptors known to absorb in the UV are not found inside chloroplasts (reviewed in 42). Second, the photoreceptors that are known to absorb in the UV-A can be excluded: cryptochromes and phototropins can be excluded based on the lack of response to blue light, and the UV-B receptor UVR8 can be excluded based on the fact that UV-B did not elicit the response and a *uvr8* mutant exhibited normal light-induced D1 synthesis. Furthermore, our conclusion that PSII damage is the trigger for D1 repair synthesis is similar to that from studies in cyanobacteria, although the response in cyanobacteria is believed to be primarily at the transcriptional level (70,71).

### Coupling of *psbA* Translation to D1 Damage and Assembly via the HCF244/OHP1/OHP2 Complex: a Working Model

Our finding that *psbA* ribosome occupancy does not decrease in the dark in *hcf136* mutants revealed a molecular link between light-regulated *psbA* translation and PSII assembly/repair. HCF136 (Ycf48 in cyanobacteria) collaborates with the HCF244/OHP1/OHP2 complex (Ycf39, HliD, HliC in cyanobacteria) to mediate early steps in assembly of the PSII reaction center (32,34,35,62,63,72,73). HCF244 and the OHPs are core subunits that interact directly with one another, whereas HCF136/Ycf48 copurified with the complex in some cases but not others (32,34,35,62,63). The HCF244/Ycf39 complex is required for both PSII assembly and repair, and has been proposed to deliver chlorophyll to nascent D1, scavenge chlorophyll from degraded D1, or protect nascent D1 from photodamage (32,34,35,63). In addition, the HCF244 complex in plants is required for *psbA* translation initiation (33,46), evoking our recent proposal that D1 autoregulates *psbA* translation via allosteric interactions within the complex (33). Results presented here allow us to expand and elaborate on this model.

The Ycf39 complex in cyanobacteria is found in two complexes that represent consecutive stages in PSII assembly: one form, called the D1 module, includes Ycf39/HliD/HliC/Ycf48 together with the PSII proteins D1 and PsbI; the second, denoted RCII*, includes, in addition, the PSII subunits D2, PsbE, and PsbF (63,74,75). A complex similar to RCII* has been detected in plants (32,34). There is evidence that HCF136/Ycf38 acts upstream of the HCF244/Ycf39 complex during PSII assembly and facilitates the incorporation of D1 into the complex (see Fig. 6 bottom) (30,31,63,76).

**Fig. 6.**
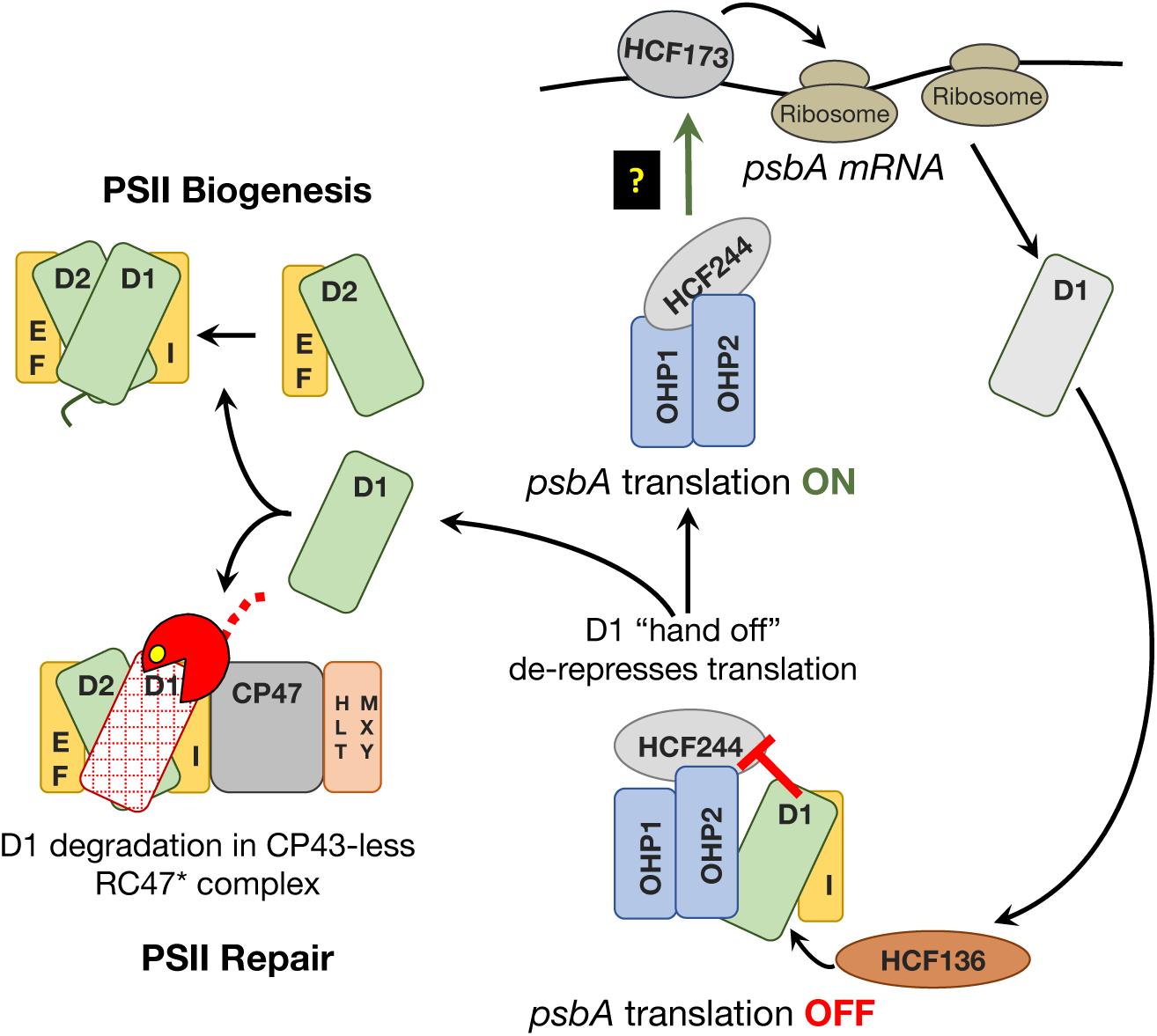
Model for the coupling of *psbA* translation to light-induced D1 damage. The HCF244/OHP1/OHP2 complex is diagrammed at bottom with bound D1 and PsbI. This form of the complex is referred to as the D1 module in *Synechocystis* (63). OHP1 and OHP2 are integral thylakoid proteins, whereas HCF244 is bound to the stromal face of the membrane (32,34,35,63). Current data suggest that HCF136 (on the luminal face of the membrane) is required to insert nascent D1 into the complex, possibly in conjunction with the insertion of chlorophyll (29-31,64). We posit that the presence of D1 in this complex inhibits the ability of the stroma-exposed components (HCF244 and OHP2) to activate *psbA* translation. Repression is relieved when D1 is transferred out of the complex, either to a repair intermediate from which damaged D1 had been removed (bottom) or to the “D2 module” during PSII biogenesis (top). We show the canonical RC47* complex (6,7) as the repair intermediate that accepts nascent D1, but the mechanism would be similar with the alternative repair intermediate detected recently in *Synechocystis* (77). We propose that the D1-less HCF244 complex communicates with HCF173 in the stroma, which is known to bind the *psbA* 5’ UTR and to activate translation initiation (48,79,81).

Consideration of our results in light of these findings leads us to propose the working model illustrated in Figure 6. According to this scheme, the presence of D1 in the HCF244/OHP1/OHP2 complex interferes with its ability to activate *psbA* translation. We propose that this inhibitory interaction is relieved by light-induced proteolysis of D1, which produces an “empty” repair intermediate to which HCF244/OHP1/OHP2 can hand off nascent D1 (Fig. 6, bottom left). This acceptor for nascent D1 could be the CP43-less reaction center complex that is generally believed to be the site of D1 degradation during PSII repair (Fig. 6 bottom left) (6,7), or it could be akin to the recently detected RC* repair intermediate detected in *Synechocystis* (77).

This model inserts a new step into current PSII repair scenarios, which posit the direct incorporation of nascent D1 into a CP43-less intermediate from which damaged D1 had been removed (6,7,14). Instead, we posit that HCF244/OHP1/OHP2, in cooperation with HCF136, are the first to engage nascent D1, and that the HCF244 complex then hands D1 off to the D1-less repair intermediate. According to this view, the HCF244 complex and HCF136 perform the same biochemical functions during both PSII repair and *de novo* PSII biogenesis. Thus, this model can readily be extended to *de novo* PSII biogenesis by positing that the transfer of D1 to the next assembly intermediate (the D2 module) will also de-repress *psbA* translation (Fig. 6 top left). This mechanism might underlie the assembly-linked regulation of D1 synthesis observed in Chlamydomonas (18), where it was proposed that translational efficiency is set to the rate of D1 assembly within a complex along the PSII biogenesis pathway. HCF244, OHP1, OHP2 are conserved (but unstudied) in Chlamydomonas, and are excellent candidates for the hub of this mechanism.

In this study, we present two new key pieces of evidence for this model. First, we observed that *psbA* ribosome occupancy remains high in the dark in *hcf136* mutants. Given that HCF136 acts upstream of the HCF244 complex during D1 assembly (29-31,64,65), the constitutively high *psbA* ribosome occupancy in *hcf136* mutants correlates with a predicted D1-less HCF244 complex under all light conditions. Second, our data strongly suggest that light-induced *psbA* ribosome recruitment and D1 synthesis are triggered by D1 damage. This strengthens evidence for the early model positing that D1 synthesis is coupled to degradation of photodamaged D1 (27,28,68). Adir and Ohad (27) observed that the rate of light-induced D1 synthesis in Chlamydomonas correlates with the rate of D1 degradation, and suggested that the damaged PSII reaction center couples light-dependent damage with D1 synthesis. Our data support this general scheme, and our model, which incorporates many advances made since that time, elaborates on the nature of the regulatory complexes and interactions.

An important aspect that is not addressed by our data concerns how the HCF244/OHP1/OHP2 complex stimulates the recruitment of ribosomes to *psbA* mRNA. Ribosome-free *psbA* mRNA is found primarily in the stroma, and *psbA* mRNA undergoing translation becomes tethered to the membrane only after D1’s first transmembrane segment emerges from the ribosome (78-80). Therefore, our model demands a means to communicate changes in the HCF244/OHP1/OHP2 complex at the thylakoid membrane to the pool of untranslated *psbA* mRNA in the stroma (black box in Fig. 6). A protein called HCF173 is a prime candidate for being involved in this membrane-stroma communication. HCF173 is required specifically for the recruitment of ribosomes to *psbA* mRNA (48,81), it partitions between the stroma and the stromal face of the thylakoid membrane (48,82), and it was pulled down with tagged OHP1 from solubilized thylakoid membranes (34). Furthermore, HCF173 associates with the 5’-untranslated region (UTR) of *psbA* mRNA (48,79,82), and it is the only protein known to bind the *psbA* mRNA and also to be important for *psbA* translation in plants. As such, our working model posits that the HCF244/OHP1/OHP2 complex impacts *psbA* translation by affecting the activity of HCF173, which in turn, regulates *psbA* ribosome recruitment by binding the 5’ UTR of the *psbA* mRNA. Clarifying the nature of the interactions among these components, and how those interactions change in response to light, D1 damage, and PSII assembly are important areas for future investigation.

## Materials and Methods

Additional information is provided in *SI Appendix, Materials and Methods*.

### Plant Material

The maize Zm*-psa3* (Zm-*psa3-1*), *pet2* (*pet2-1/-3*), Zm-*hcf107*, and Zm-*hcf136* mutants were described previously (33,50,58,59). The maize PAB ortholog (Zm-PAB) is encoded by Zm00001d048524 (B73 RefGen_v4). Evidence for its orthology with Arabidopsis *pab* (AT4G34090) (49) can be found at http://cas-pogs.uoregon.edu/#/pog/14960. Two Zm-*pab* transposon insertion alleles were recovered from the Photosynthetic Mutant Library collection (83) (described in *SI Appendix, Fig. S1A*); the heteroallelic progeny of a complementation test cross were used for the experiments here.

The Arabidopsis *psa3* (SAIL_503_B01) and *hcf107-3* (SALK_079285C) mutants were described previously (50,81). The Arabidopsis *stn7-1/stn8-1* double mutant was a gift of Sacha Baginsky (Martin Luther University) and was described in ref (55). Arabidopsis *hcf136* was obtained from Jörg Meurer, Susanne Paradies, and Peter Westhoff, and is the same allele studied previously (29,33). Arabidopsis *hcf208* was obtained from Peter Westhoff and was described in ref (47). Arabidopsis *uvr8-6* was obtained from Roman Ulm and Emilie DeMarsy, and was described in ref (84). See *SI Appendix, Materials and Methods* for additional information.

### Action Spectrum Experiment

Arabidopsis Col-0 seedlings were grown for 18-21 days on MS medium and were then acclimated to 8 μmol m^−2^ s^−1^ (fluorescent bulb Sylvania F40/D41/SS) for 30 min at midday. Supplemental lighting was then provided for 15 minutes, with the lids of the plates removed. Leaves were then harvested for ribo-seq or were excised and used for *in vivo* pulse labeling in the same light condition. See *SI Appendix, Materials and Methods* for additional information.

### DCMU, Nigericin, and CCCP Treatments

Arabidopsis Col-0 seedlings were grown on MS plates as described for the action spectrum experiment. Wounds were created by excising cotyledons. Each seedling with attached roots was transferred to 0.5 ml MS medium (2% (w/v) sucrose, pH 5.7) containing drug in a well of a clear 24-well plate. DCMU (10 mM stock in 50% ethanol), nigericin (5 mM stock in 95% ethanol) and CCCP (carbonyl cyanide 3-chlorophenylhydrazone, 100 mM stock in DMSO) were diluted in MS medium to final concentrations of 20 µM, 25 µM, and 100 µM, respectively. Mock treatments with the solvent were used as controls.

The DCMU treatments used 14-day-old Col-0 seedlings grown under long day conditions (see above). The seedlings were incubated in the DCMU solution in darkness for 2 h. One set of seedlings was then shifted to light (8 μmol m^−2^ s^−1^) and another was maintained in darkness. After 15 min in light, the tissues were harvested for ribo-seq or the leaves were excised and used for *in vivo* pulse labeling in buffer containing DCMU. The nigericin and CCCP experiments used 23-day-old Col-0 seedlings grown in short days. The seedlings were incubated in nigericin or CCCP solutions for 15 min in the light (8 μmol m^−2^ s^−1^). The leaves were then excised for in *vivo* pulse labeling in buffer containing nigericin or CCCP.

### *In Vivo* Pulse Labeling

*In vivo* labeling of maize and Arabidopsis seedlings was performed as described previously (19), with minor modifications. Unless otherwise indicated in figures, seedlings were acclimated to the assay light for 30 min prior to the experiment. Briefly, maize seedlings were radiolabeled for 15 minutes by applying ^35^S-methionine/cysteine to a leaf abrasion, whereas excised Arabidopsis leaves were submerged in labeling mix for 20 minutes. The “1-h dark” labeling was performed from 40 to 60 min, or 45-60 min after the shift to dark in Arabidopsis and maize, respectively. The reillumination labeling was performed starting 15 minutes after the shift back to light. Leaf extracts were separated by SDS-PAGE, transferred to nitrocellulose and imaged with a Storm Phosphorimager (GE Healthcare). Gel loading was normalized based on radiolabeled cytosolic proteins in maize, and based on either chlorophyll or total leaf protein mass in Arabidopsis (as stated in figure legends). Additional details are provided in *SI Appendix, Materials and Methods*.

### Ribo-seq and RNA-seq

Maize ribosome footprints were prepared from the apical portion of the second and third leaves to emerge, excising above the ligule of leaf 1. One seedling was used per replicate, except for the first replicate of Zm-*hcf107*, which used two pooled seedlings. Arabidopsis ribosome footprints were prepared from the aerial portions of pools of three to five seedlings. Tissue was flash-frozen in liquid N_2_ and stored at -80°C until used. Ribosome footprint preparation, total RNA extraction, sequencing library construction and data analysis were performed as described previously (19), with minor modifications described in *SI Appendix Materials and Methods*.

### Immunoblot Analysis and Antibodies

Immunoblots were performed as described previously (85). Antibodies to AtpB, D1, PsaD and PetD were generated by our group and were described previously (86). The antibody to D2 was obtained from Agrisera.

## Data Availability Statement

All data obtained for this study are presented within the main text, *SI Appendix* and *Dataset S1*. Illumina sequencing data have been deposited in the NCBI Sequence Read Archive under accession PRJNA612079.

## Acknowledgments

We are grateful to Susan Belcher and Roz Williams-Carrier for expert technical assistance, Helmut Kirchhoff, Ryoichi Tanaka, Josef Komenda, and Roman Sobotka for helpful discussions, Amy Turner for loaning spectrometers, and Nick Stiffler for assistance with data deposition. We also appreciate the generous gifts of mutant Arabidopsis lines from Sacha Baginsky, Jörg Meurer, Susanne Paradies, Peter Westhoff, Emilie Demarsy, and Roman Ulm. This research was funded by the US National Science Foundation, grant numbers MCB-1616016 and IOS-1339130.

## SI Appendix

### SI Materials and Methods

#### Plant Material

Maize seedlings were grown under cycles of 16-h light at 28°C and 8-h dark at 26°C with light intensity of 250-350 μmol m^−2^ s^−1^ (mixture of fluorescent Sylvania F72T12/CW/VHO or F48T12/CW/HO with incandescent Sylvania 100A/DLSW/VERT/RP). Phenotypically wild-type siblings were used as the wild-type controls in each experiment. Arabidopsis seeds were grown on 1x MS medium supplemented with 2 or 3% (w/v) sucrose and 0.3% (w/v) Phytagel, pH 5.7. Plants used for the *hcf107, hcf136*, action spectrum, nigericin, and CCCP treatments were grown in short days (10-h light/14-h dark) at 22°C. Plants used for the At-*psa3, hcf208*, and DCMU experiments were grown in long days (16-h light/8-h dark) at 22 °C. A light intensity of 80-100 μmol m^−2^ s^−1^ (fluorescent Sylvania F72T12/CW/VHO or F48T12/CW/HO) was used for Arabidopsis growth, except in the case of the *hcf208* line, which was grown under lower light intensity (30-50 μmol m^−2^ s^−1^) to increase PSII abundance. Phenotypically normal siblings were used as wild-type controls except in the *stn7-stn8* experiment, which used the Col-0 progenitor as the wild-type.

#### Action Spectrum Experiment

Leaves from four seedlings were pooled for each ribo-seq sample and leaves from two or three seedlings were pooled for pulse labeling. The following sources were used for supplemental lighting: 25 μmol m^−2^ s^−1^ white light (incandescent bulb Sylvania 100A/DLSW/VERT/RP), 5 μmol m^−2^ s^−1^ UV-B (M-26XV Benchtop Variable Transilluminator, UVP), 5 μmol m^−2^ s^−1^ UV-A (UVL-56 Handheld UV Lamp, UVP), 5 μmol m^−2^ s^−1^ blue, green or red light emitting diodes (LEDs) (Remix Headlamps, Princeton Tec) with maximum emission wavelengths (λ) of 314, 370, 469, 522 and 635 nm, respectively. The spectra of UV-A and UV-B lights were determined using a CCS200 Compact Spectrometer (Thorlabs) from 195 to 1020 nm and FLAME-S-UV-VIS-ES Spectrometer (Ocean Optics) from 180 to 880 nm, respectively. The spectra of LEDs were determined with a CCS200 Compact Spectrometer (Thorlabs) from 195 to 1020 nm and an ILT350 Illuminance Spectrometer (International Light Technology) from 380 to 780 nm. The intensities of LEDs with wavelengths within the photosynthetic active radiation (PAR) range were determined using a MQ-200 quantum meter (Apogee Instruments). To determine the intensities of UV-A and UV-B, the light energy flux (irradiance in W cm^−2^) was measured using an ILT1700 Research Radiometer (International Light Technology). The light photon energy (E_p_) was calculated using the maximum emission wavelength from E_p_ = *hc / λ* where *h* is Planck’s constant and *c* is speed of light. The light intensity (μmol photon m^−2^ s^−1^) is the irradiance I / (E_p_ x Avogadro constant).

#### *In Vivo* Pulse Labeling

Details for specific experiments were as follows. A light intensity of 8 μmol m^−2^ s^−1^ (Sylvania F40/D41/SS) was used for the drug treatments. A light intensity of 100 μmol m^−2^ s^−^ 1 (AgroMax F24/T5/HO) was used for mutant assays. Light conditions for the action spectrum analyses are indicated in figures. Unless otherwise indicated, rosette leaves from two to three seedlings (approximately two weeks post germination, 6-8 leaves per labeling) were excised and placed in a clear 24-well plastic plate containing 135 μl of labeling buffer (100 μg/mL cycloheximide, 1x MS medium, 2% (w/v) sucrose, pH 5.7). 15 μl of EasyTag Express ^35^S Protein Labeling Mix was added immediately to initiate labeling. For *uvr8*, the young leaves from 55-day-old plants were used for labeling (total of 6-8 leaves pooled from 2 plants per labeling). For the *stn7-stn8* mutant, leaves were pre-incubated for 30 min in labeling buffer containing 25 μg/mL cycloheximide prior to addition of the radiolabeled amino acids. After the 20 min labeling period, leaves were washed twice in labeling buffer lacking radiolabeled amino acids and frozen in liquid nitrogen. The homogenization buffer was as described previously (1), with 5 mM MgCl_2_ added for the At-*psa3* and At-*hcf208* experiments to improve recovery of thylakoid membranes in the pellet. The membrane fraction was collected by centrifugation at 10,000 x g for 5 min at 4°C and washed twice. Membrane fractions were resolved on 11% polyacrylamide SDS-PAGE gels containing 6 M urea, whereas total leaf extracts were resolved on 4-20% polyacrylamide SDS-PAGE gels.

#### Ribo-seq and RNA-seq

Ribosome footprints were prepared as described previously (1). In brief, RNase I was used to generate monosomes, monosomes were pelleted through a sucrose cushion, RNA was extracted from the pelleted material, RNA between 20 and 40 nucleotides was gel purified, and the gel-purified RNA was used as input for the NEXTflex Small RNA Sequencing Kit v3 (Perkin Elmer). rDNA depletion was performed using species-specific sets of biotinylated DNA oligonucleotides (1). Aliquots of the same tissue homogenates were used for RNA-seq, which used the Ribo-Zero rRNA Removal Kit (Plant Leaf) (Illumina) and the NEXTflex qRNA-Seq Kit v2 (Perkin Elmer). Libraries were sequenced on a HiSeq 4000 instrument (Illumina), with read lengths of 75 or 100 nucleotides.

Sequencing data were processed as described in (2) with the following modifications. Trimmed reads were sequentially aligned to the chloroplast, mitochondrial and nuclear genomes using STAR (3) instead of Bowtie 2. Maize reads were aligned to GenBank accession X86563 (chloroplast genome) and B73 RefGen_v4 assembly version 38 (maizegdb.org) (nucleus and mitochondria). Arabidopsis reads were aligned to the TAIR10 genome and annotation (arabidopsis.org). Read counting was performed using featureCounts (4) instead of the in-house scripts used previously. Read counts for chloroplast genes excluded the first 10 nucleotides of each ORF to avoid counting the ribosome pileup at the start codon. Chloroplast read counts were normalized by using reads per kilobase of coding sequence (CDS) per million reads mapped to all chloroplast CDSs (cpRPKM). Read mapping statistics and cpRPKM values from all the experiments are provided in *SI Appendix*, Dataset S1.

### SI Figures

**Fig. S1.**
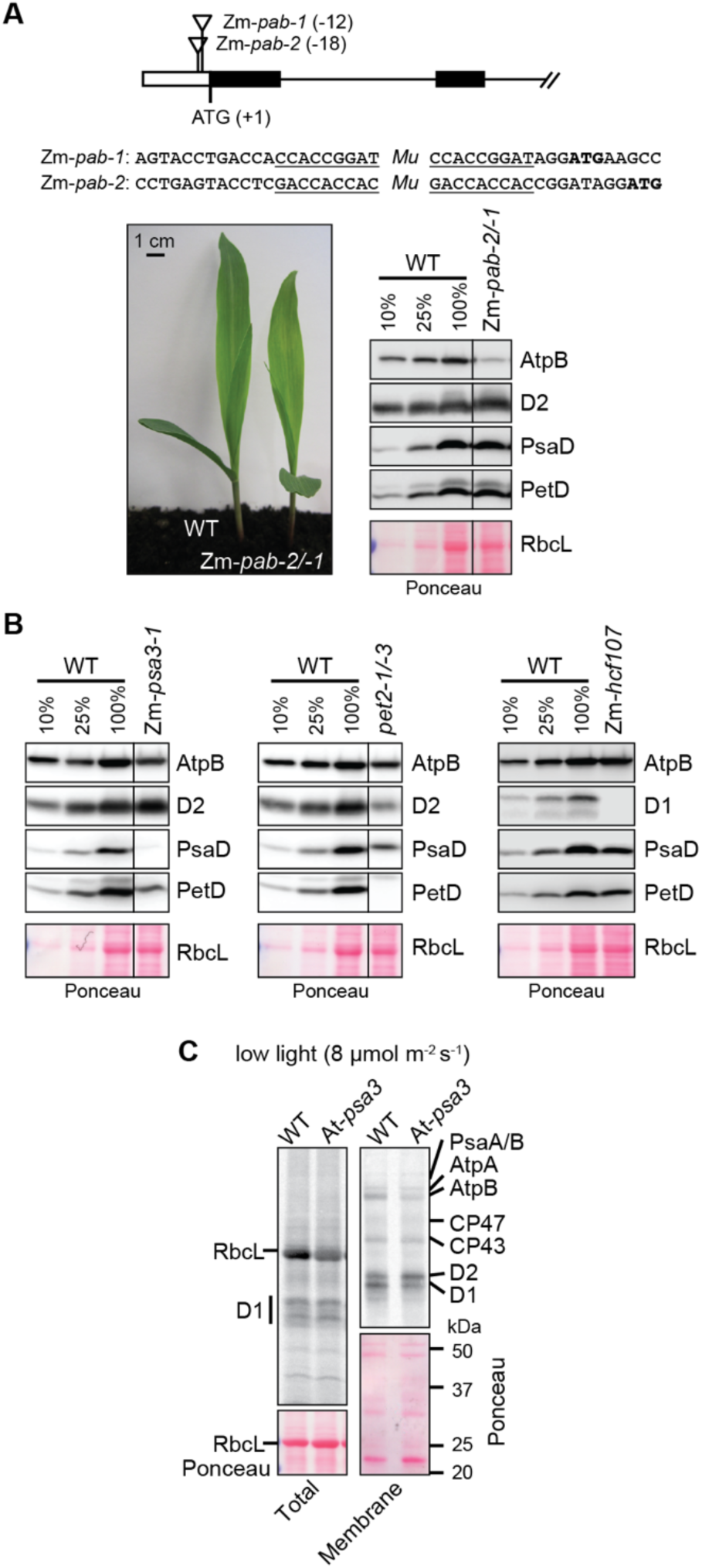
Additional information about mutants analyzed in Figures 3, 4, and 5. (*A*) Maize *pab* mutants. Two transposon insertion alleles were recovered from the Photosynthetic Mutant Library (5). The position of the insertions with respect to the start codon is diagrammed, and the sequences of the insertion sites are shown with the target site duplications underlined. The heteroallelic progeny of a complementation test cross (Zm-*pab-2/-1*) were used for the experiments here. The phenotype of seedlings at the stage used here (grown for 8 days in soil) is shown. The immunoblots to the right show the abundance of representative core subunits of photosynthetic complexes. Replicate blots were probed to detect AtpB (subunit of the chloroplast ATP synthase), D2 (subunit of PSII), PsaD (subunit of PSI), and PetD (subunit of the cytochrome *b*_*6*_*f* complex). An excerpt of one of the Ponceau S-stained blots illustrates relative sample loading and the abundance of the large subunit of Rubisco (RbcL). The specific loss of AtpB is as expected based on analysis of the orthologous mutant in Arabidopsis (6). Lines separate non-contiguous lanes from the same exposure of the same blot. (*B*) Immunoblot validation of maize mutants used here that were reported previously (7-9). Lines separates non-contiguous lanes from the same exposure of the same blot. (*C*) Pulse labeling of At-*psa3* in low intensity light (8 μmol m^−2^ s^−1^). Labeling was performed in the presence of cycloheximide for 20 min. Total leaf lysates (left) and membrane fractions (right) were resolved by SDS-PAGE and transferred to nitrocellulose, and radiolabeled proteins were detected by phosphorimaging. The imaged blots were also stained with Ponceau S to illustrate relative sample loading.

**Fig. S2.**
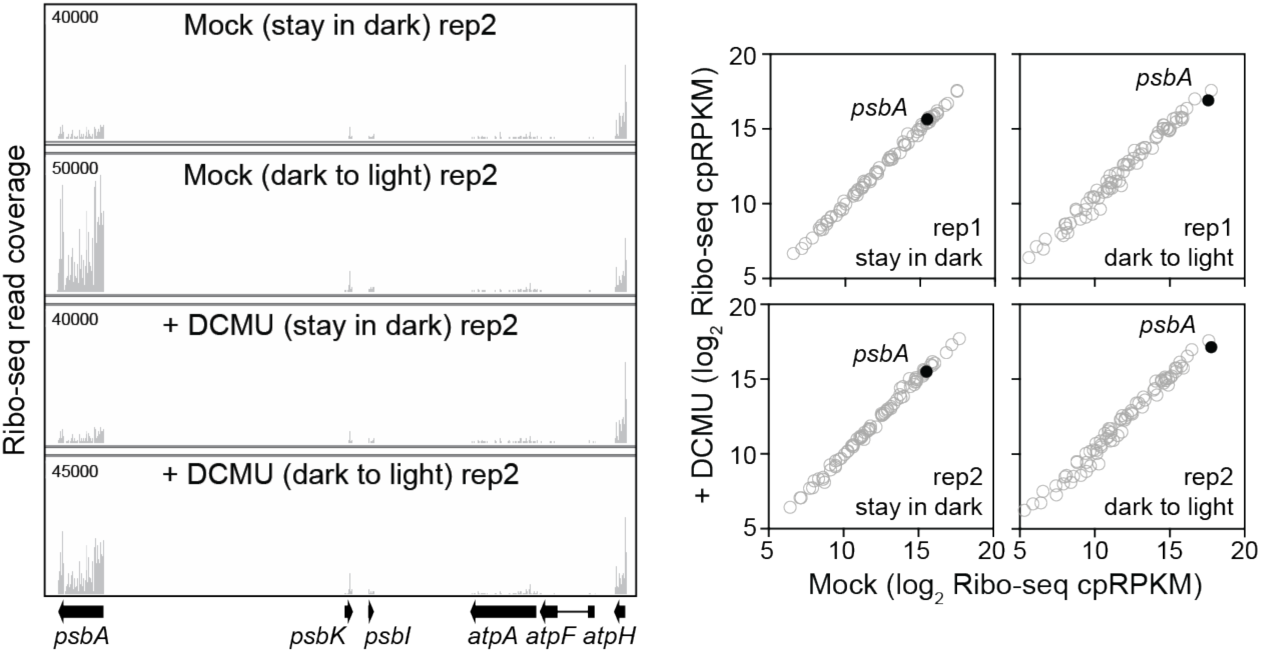
Additional ribo-seq data to support the DCMU experiment shown in Figure 5*A*. Arabidopsis (Col-0) was treated with DCMU as shown in Fig. 4*C*. The left panel shows Integrated Genome Viewer (IGV) screen captures from a replicate of the ribo-seq experiment shown in Fig. 5*A*. Right panels show scatter plot comparisons of ribosome footprint abundance for all chloroplast genes in the DCMU-treated and untreated (mock) samples. Each symbol represents one gene. Values are expressed as reads per kilobase in the ORF per million reads mapped to chloroplast ORFs (cpRPKM).

**Fig. S3.**
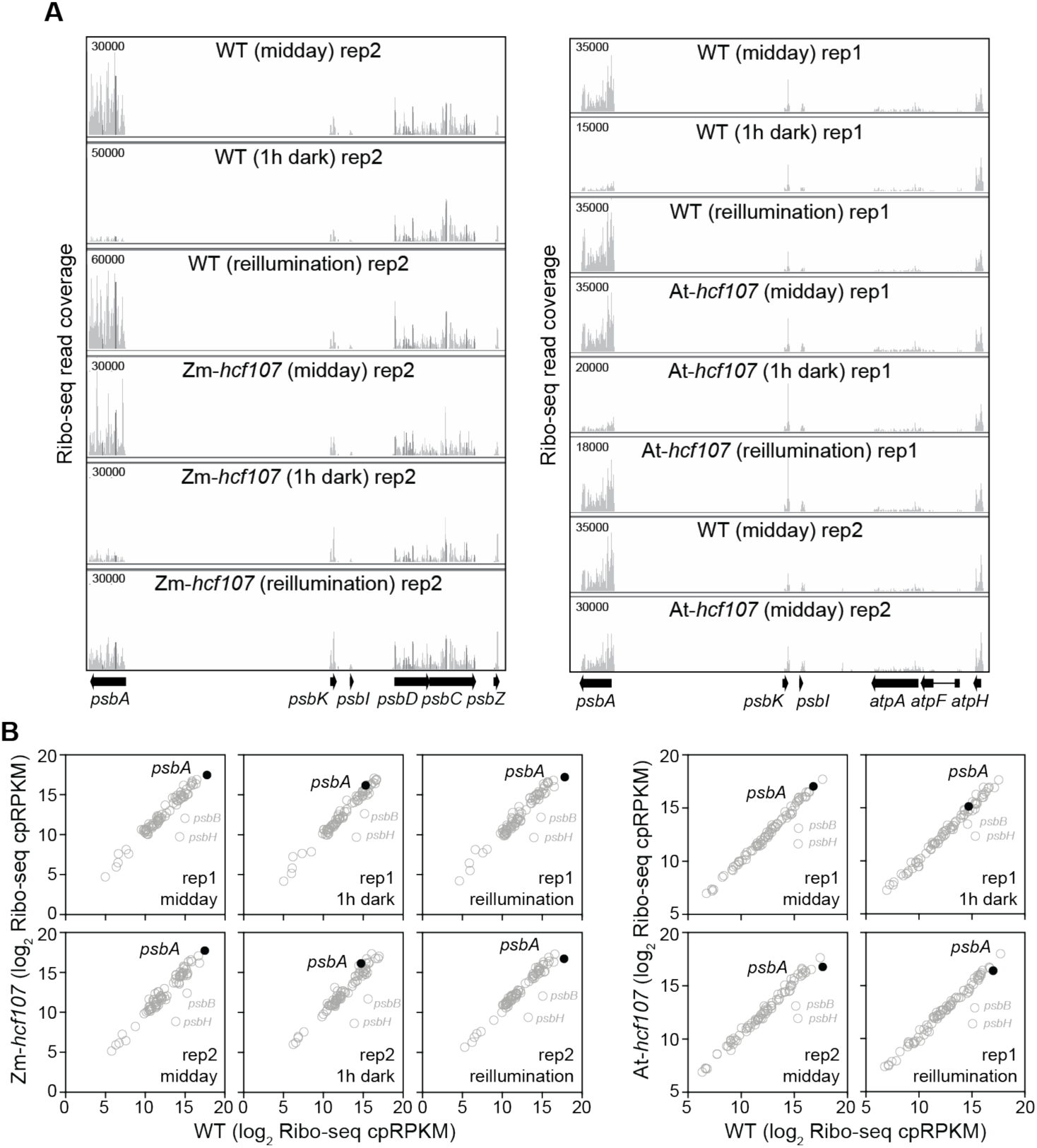
Additional data to support the ribo-seq analysis of At-*hcf107* and Zm-*hcf107* mutants shown in Figure 5*B*. The experiments were performed as described in Fig. 5*B*. *(A)* Screen captures from the Integrated Genome Viewer (IGV) showing the distribution of ribosome footprints along *psbA* and adjacent ORFs. The maximum Y-axis values are shown in the upper left, and were chosen such that coverage of adjacent ORFs were similar. *(B)* Scatter plots comparing ribosome footprint abundance for all chloroplast genes in the mutant and its phenotypically normal siblings (WT). Each symbol represents one gene. Values are expressed as reads per kilobase in the ORF per million reads mapped to chloroplast ORFs (cpRPKM). The second replicate of the Arabidopsis midday analysis was published previously (8).

**Fig. S4.**
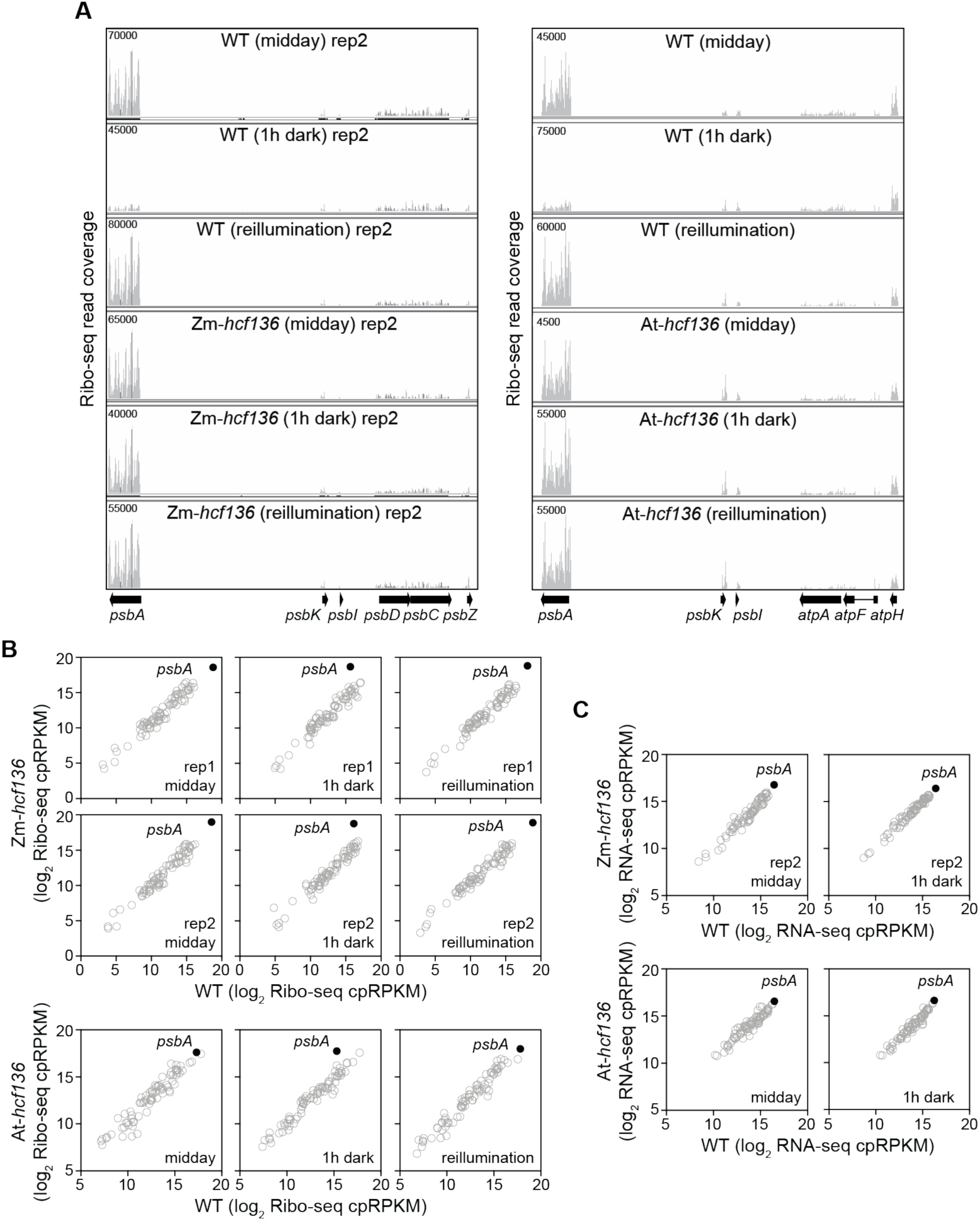
Additional data to support the ribo-seq analysis of At-*hcf136* and Zm-*hcf136* mutants shown in Figure 5*C*. The results from the midday samples were published previously (10), but these were analyzed in parallel with the corresponding 1-h dark and 15-min reillumination samples. (*A*) Screen captures from the Integrated Genome Viewer (IGV) showing the distribution of ribosome footprints along *psbA* and adjacent ORFs. The maximum Y-axis values are shown in the upper left, and were chosen such that coverage of adjacent ORFs were similar. *(B)* Scatter plots comparing ribosome footprint abundance for all chloroplast genes in the mutant and its phenotypically normal siblings (WT). Each symbol represents one gene. Values are expressed as reads per kilobase in the ORF per million reads mapped to chloroplast ORFs (cpRPKM). *(C)* Scatter plots comparing mRNA abundance (RNA-seq) for all chloroplast genes in the mutant and its phenotypically normal siblings (WT). RNA was prepared from the same lysates used for ribo-seq. Each symbol represents one gene. Values are expressed as reads per kilobase in the ORF per million reads mapped to chloroplast ORFs (cpRPKM).

